# Comparative biochemistry of four polyester (PET) hydrolases

**DOI:** 10.1101/2020.11.20.392019

**Authors:** Jenny Arnling Bååth, Kim Borch, Kenneth Jensen, Jesper Brask, Peter Westh

## Abstract

The potential of bioprocessing in a circular plastic economy has strongly stimulated research in enzymatic degradation of different synthetic resins. Particular interest has been devoted to the commonly used polyester, poly(ethylene terephthalate) (PET), and a number of PET hydrolases have been described. However, a kinetic framework for comparisons of PET hydrolases (or other plastic degrading enzymes) acting on the insoluble substrate, has not been established. Here, we propose such a framework and test it against kinetic measurements on four PET hydrolases. The analysis provided values of k_cat_ and K_M_, as well as an apparent specificity constant in the conventional units of M^−1^s^−1^. These parameters, together with experimental values for the number of enzyme attack sites on the PET surface, enabled comparative analyses. We found that the PET hydrolase from *Ideonella sakaiensis* was the most efficient enzyme at ambient conditions, and that this relied on a high k_cat_ rather than a low K_M_. Moreover, both soluble and insoluble PET fragments were consistently hydrolyzed much faster than intact PET. This suggests that interactions between polymer strands slow down PET degradation, while the chemical steps of catalysis and the low accessibility associated with solid substrate were less important for the overall rate. Finally, the investigated enzymes showed a remarkable substrate affinity, and reached half the saturation rate on PET, when the concentration of attack sites in the suspension was only about 50 nM. We propose that this is linked to nonspecific adsorption, which promotes the nearness of enzyme and attack sites.

## Introduction

Polyethylene terephthalate (PET) is a copolymer of terephthalic acid (TPA) and ethylene glycol (EG) linked by ester bonds (Fig. 1), and among the most produced plastics. The stable nature of the PET molecule has been an advantage for its use in fibers, films and packaging, but this characteristic is also becoming an increasing problem, as it leads to accumulation of PET in the environment (1–3). This causes a growing need for cost-effective means to recycle PET waste for example through monomer recovery and re-synthesis of virgin residue. Recent progress suggests that this may be achieved through bioprocessing (1,4), and one crucial requirement for this is the development of efficient enzymes for PET degradation.

**Figure 1.**
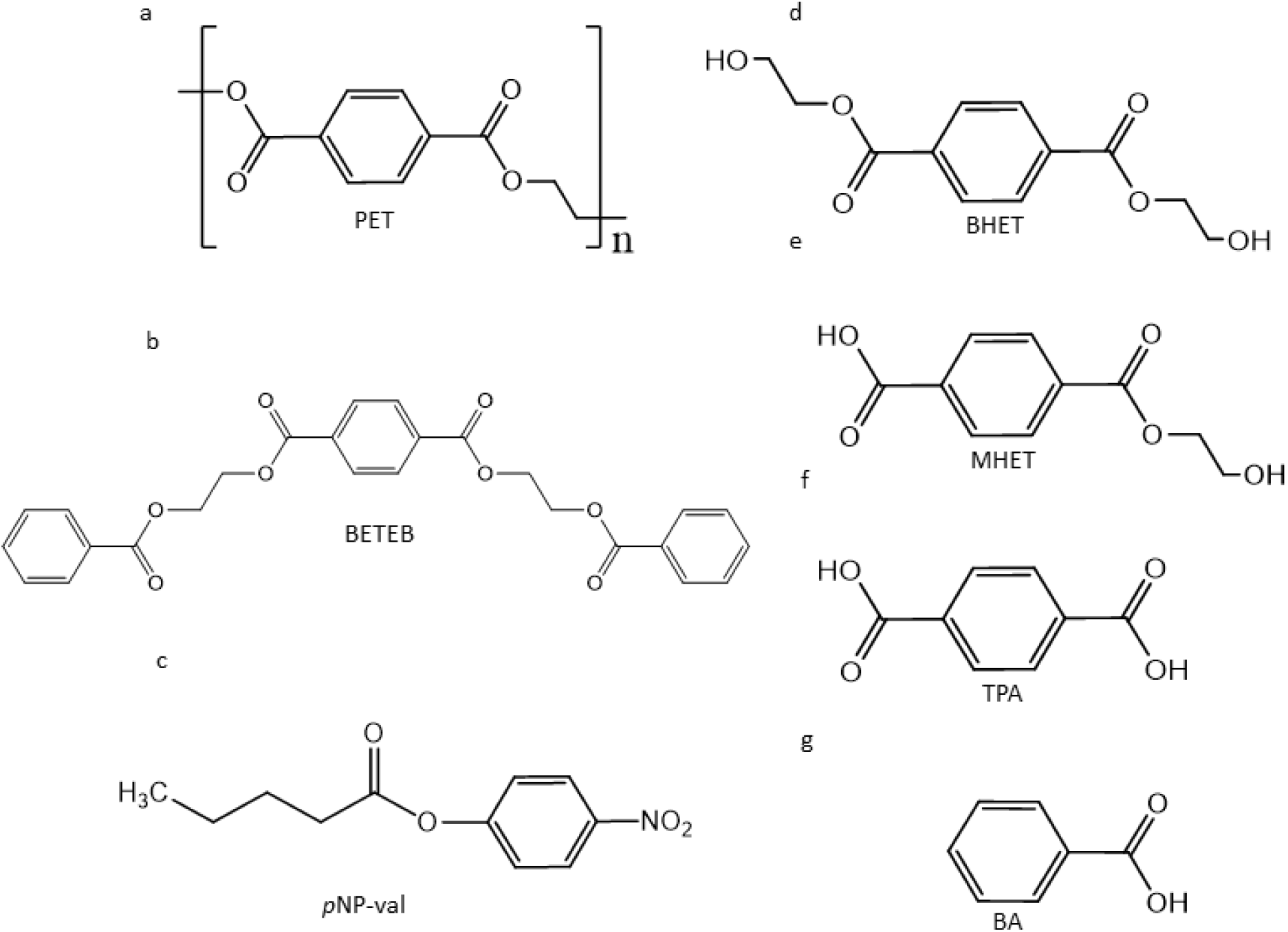
Substrates and products discussed in this study. a) Chemical structure of the intact PET polymer; b) In-house synthesized PET fragment, BETEB, with three aromatic rings; c) Conventional substrate analog, *p*NP-val, for cutinases; d) PET fragment, BHET, with two ester bonds; e) The repeating unit, MHET, of the PET polymer; f) The unesterified diacid, TPA; and g) monoacid BA. BHET and MHET were both observed as reaction products from PET hydrolysis, but were additionally used as substrates. BA was a reaction product from BETEB hydrolysis reactions. TPA was observed as a minor reaction product from PET, BETEB, BHET and MHET hydrolysis and is the constituent monomer of the PET polymer.

Research on enzymatic degradation of PET has been ongoing for over 20 years, and enzymes classified as cutinases have been described as the most promising PET hydrolases (2,3,5). Cutinases are typical serine hydrolases, with an α/β fold and a catalytic triad consisting of serine, histidine and aspartate (6). Compared to lipases and many ester-active enzymes, cutinases exhibit a flat, surface exposed active site, and this have been described as essential for their interaction with the PET polymer, which is bulkier than their preferred, aliphatic substrate, cutin (7,8). PET hydrolases have been described as enzymes with low to moderate turnover rates, reflecting the fact that PET is an unnatural substrate for these enzymes (2). However, in 2016 the bacterium *Ideonella sakaiensis* was discovered, possessing an unprecedented capacity to use PET as a source of carbon and energy. This bacterium was isolated from a PET-rich environment and secretes a PET hydrolase, which is homologous to cutinases, and may represent a short evolutionary adaptation to the synthetic substrate. Due to its superior PET degrading ability and a significantly lower activity on natural, aliphatic polyesters it has been categorized into a novel family of enzymes, named PETases (EC 3.1.1.101) (5). The discovery of the *I. sakaiensis* PETase has led to several structural studies, which have improved our understanding of the catalytic process and consequently outlined strategies for rational engineering of enzyme variants with improved activity against the synthetic substrate (9–12). However, this progress in structural understanding has not been paralleled by biochemical investigations. Thus, while some studies have reported kinetic parameters for PET hydrolases (4,8,13–17), most functional assessments have used long contact times and empirical discussions of either end-point concentrations or progress curves (18–23). Results from the latter type of work primarily elucidate time- and dose requirements to achieve a significant degree of polymer conversion, and are hence important for technical applications of PET hydrolases. However, in the absence of physically meaningful kinetic parameters, one can typically only make superficial analyses of structure-function relationships. This makes it difficult to compare results across different studies, and impedes the potential for interpretations regarding molecular mechanisms of the catalytic process.

The scarcity of kinetic parameters can be linked to the interfacial nature of the reaction, which gives rise to some complications that are unknown in conventional (bulk) enzyme kinetics. One particular difficulty arises because the molar concentration of the substrate is unknown. This challenges the use of mass-action kinetics and hence the conventional Michaelis-Menten (^conv^MM) framework (8,24). However, mass-action kinetics is widely used for non-biochemical, interfacial catalysis (25,26), and the normal way to handle the solid material in rate equations is to define a number of sites on the surface, which are competent for the process in question. Along these lines, we define an attack site on the PET surface as a locus, where the PET hydrolase is able to form a productive substrate complex (Fig. 2). If the density of attack sites, Γ_attack_ in units of mol sites per gram PET, can be experimentally established, it is possible to convert a substrate load (in g/L) into a molar concentration of sites (27). This is only an apparent molar concentration, because the surface sites are not evenly distributed in the suspension, but the approach opens up for the use of mass-action kinetics, and hence the derivation of rigorous kinetic parameters for a heterogeneous enzyme reaction.

**Figure 2.**
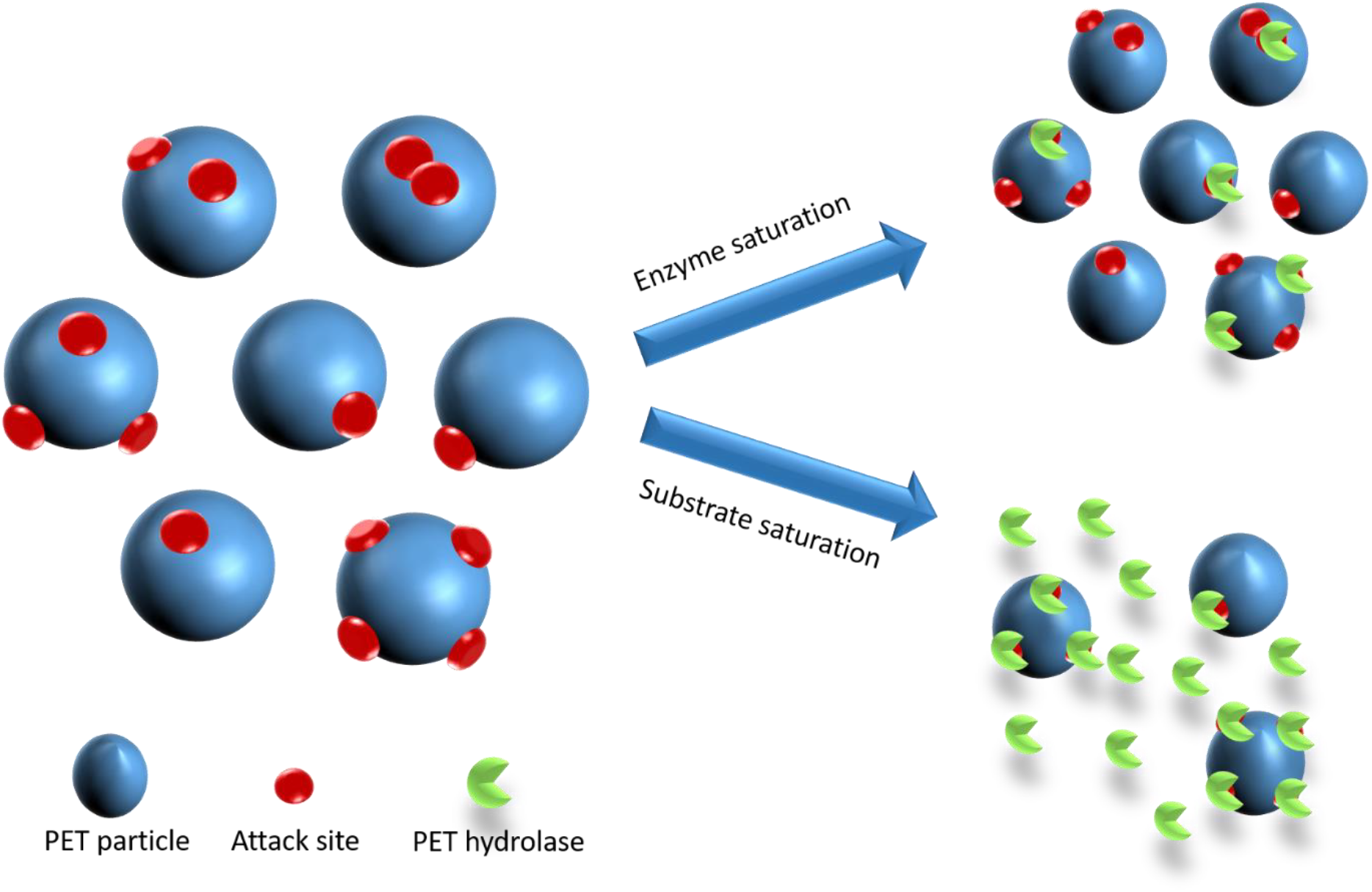
Schematic illustration of saturation under the two experimental conditions investigated here. The well-known, conventional Michaelis-Menten (^conv^MM) approach, uses an excess of substrate and initial rate measurements at a number of substrate loads. At high substrate loads, this leads to “enzyme saturation” where all enzyme is in a bound state. Conversely, the inverse (^inv^MM) approach measures initial rates at a low substrate load and gradually increasing enzyme concentrations. This leads to “substrate saturation”, where all attack sites on the substrate surface are occupied.

In the current work, we have tested a kinetic approach based on these ideas for PET hydrolases. Specifically, we analyzed rate measurements obtained under two different experimental conditions. One set of data was made in the usual limit of substrate excess, while the other was made under condition of enzyme excess. For interfacial enzyme reactions, the steady-state approximation may be justified for both of these conditions (28), and the latter, so-called inverse Michaelis-Menten (^inv^MM) framework, has occasionally been used for solid substrates (29–31) including PET (8,14,15). Both ^conv^MM plots (rate vs. substrate load) and invMM plots (rate vs. enzyme concentration) may lead to hyperbolic curves (27), and the benefit of combining these two approaches may be illustrated by considering the conditions at saturation. For convMM, saturation reflects the well-known situation, where all enzyme are engaged in a complex, and the rate becomes (*c.f.* Fig. 2)

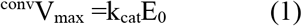

For ^inv^MM, saturation occurs at high enzyme concentration as all attack sites on the surface become occupied (and additional enzyme accumulates in the bulk). If we assume that the conversion of these sites is governed by k_cat_, the inverse saturation rate may be written

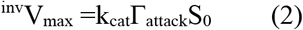

where S_0_ is the (known) mass load of substrate. The expression in eq. (2) emerges intuitively as illustrated in Fig. 2, but has been derived rigorously elsewhere (27). As ^conv^V_max_ and ^inv^V_max_ can be derived from experiments, we can estimate Γ_attack_ as the ratio of the maximal specific rates. Specifically, combining eqs. (1) and (2) yields

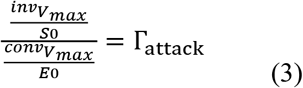

We have used these ideas to characterize a group of enzymes consisting of a catalytically improved variant of *I. sakaiensis* PETase (12), cutinases from respectively the fungus *Humicola insolens* and the bacterium *Thermobifida fusca* and a carboxyl-esterase from *Bacillus subtilis*. These enzymes have previously been reported to hydrolyze PET (8,14,21) and here we conduct a comparative kinetic analysis with respect to their activity on both polymeric PET and smaller (soluble or insoluble) model substrates, primarily PET fragments. We also report a conspicuous effect of the nonionic surfactant n-dodecyl β-D-maltoside (DDM) on the kinetic parameters of these enzymes. The results provide novel insights into the enzymatic degradation of PET, and we hope that the suggested kinetic approach may serve as inspiration for further developments of comparative approaches within this rapidly growing and technically important area of enzymology.

## Results

We measured hydrolytic rates of two cutinases (HiC and TfC), one PETase (IsP) and one carboxyl-esterase (BsCE) on both polymeric PET and a number of small ester substrates and PET fragments (see Fig. 1 for substrate structures). In addition, we studied the influence of the nonionic surfactant DDM on the enzyme kinetics. The hydrolysis of PET was quantified by the increment of UV absorption in the supernatant detected in a plate reader, while activity on smaller substrates required RP-HPLC-based detection. The plate reader-based detection method relies on assumptions discussed elsewhere, but is feasible for comparative studies (32). We made ^conv^MM analysis for all substrates. Specifically, reaction rates were plotted as a function of the load or concentration of substrate and analyzed with respect to the ^conv^MM equation, eq. (4).

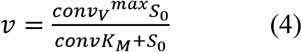

For the insoluble substrates PET and BETEB, we also conducted experiments with enzyme excess. In these cases, the reaction rates were plotted as a function of the enzyme concentration and analyzed with respect to the ^inv^MM equation, eq. (5) (27)

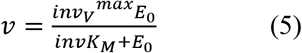

### Steady-state kinetics with PET as substrate

Three of the investigated enzymes (HiC, TfC and IsP) showed distinct activity on suspended PET powder, while no product release was detected for BsCE after 24 h incubation at 50 °C. BsCE has previously been described as an enzyme that acts on PET films, but this was based on experiments with 120 h contact time and over ten-fold higher enzyme concentration than in the current study (21). Hence, we conclude that BsCE performs poorly on polymeric PET compared to HiC, TfC and IsP. In order to decide on suitable contact times for activity measurements for the three PET-active enzymes, we made progress curves with excess of either PET or enzyme (Fig. S2). Incubation times within the linear range of these progress curves were deemed appropriate for determination of steady-state rates. The three PET-active enzymes were investigated at different temperatures below the midpoint of thermal denaturation, T_m_. At pH 8, T_m_ was 66 °C for HiC, 80 °C for TfC, 55 °C for IsP and 56 °C for BsCE.

Representative rate measurements are illustrated as conventional-(Fig. 3) and inverse-(Fig. 4) MM plots, and parameters derived by fitting respectively eq. (4) or eq. (5) are listed in Table 1 (^conv^MM) and Table 2 (^inv^MM). The two equations generally accounted well for the data except in the case of ^inv^MM for IsP. As shown in Fig. 4, this system showed the expected behavior at low enzyme concentrations, but declining rates at higher enzyme dosages. A similar behavior has been observed previously both for another cutinase hydrolyzing PET (17) and for enzymes hydrolyzing the natural polyester poly[(R)-3-hydroxybutyrate] (PHB), and this may reflect surface denaturation as the enzyme coverage increases (33–35). This correlation of high surface coverage and denaturation is widely observed for adsorbed populations of marginally stable proteins (36).

**Figure 3.**
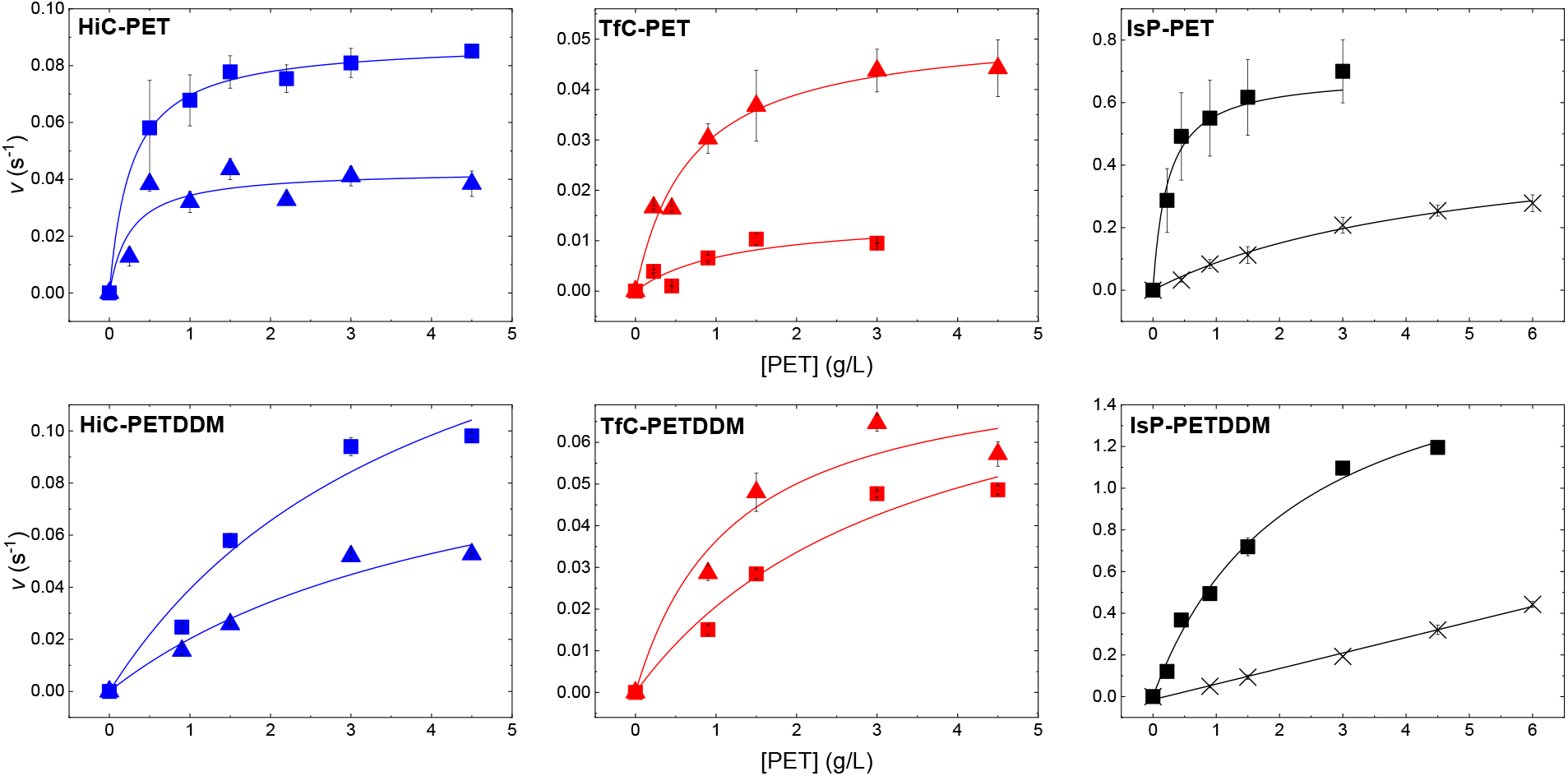
Conventional Michaelis Menten (^Conv^MM) plots for HiC (blue), TfC (red) and IsP (black). The curves show hydrolysis rates as a function of PET load in g/L. Symbols are experimental data from 2 h (IsP) or 5 h (HiC and TfC) reactions at 40 °C (crosses), 50 °C (squares) or 60 °C (triangles) with 0.03 μM enzyme. Upper panels are without the addition of the nonionic surfactant n-Dodecyl β-D-maltoside (DDM). Lower panels are with 0.0025% (w/V) DDM. Each enzyme was investigated at two experimental temperatures below their T_m_. Error bars represent standard deviations of triplicate measurements and lines are the best fit of eq. (4). For IsP-PET with DDM at 40 °C, K_M_ was much larger than the highest substrate load and these data are fitted to a straight line. Parameters derived from these results may be found in Table 1.

**Figure 4.**
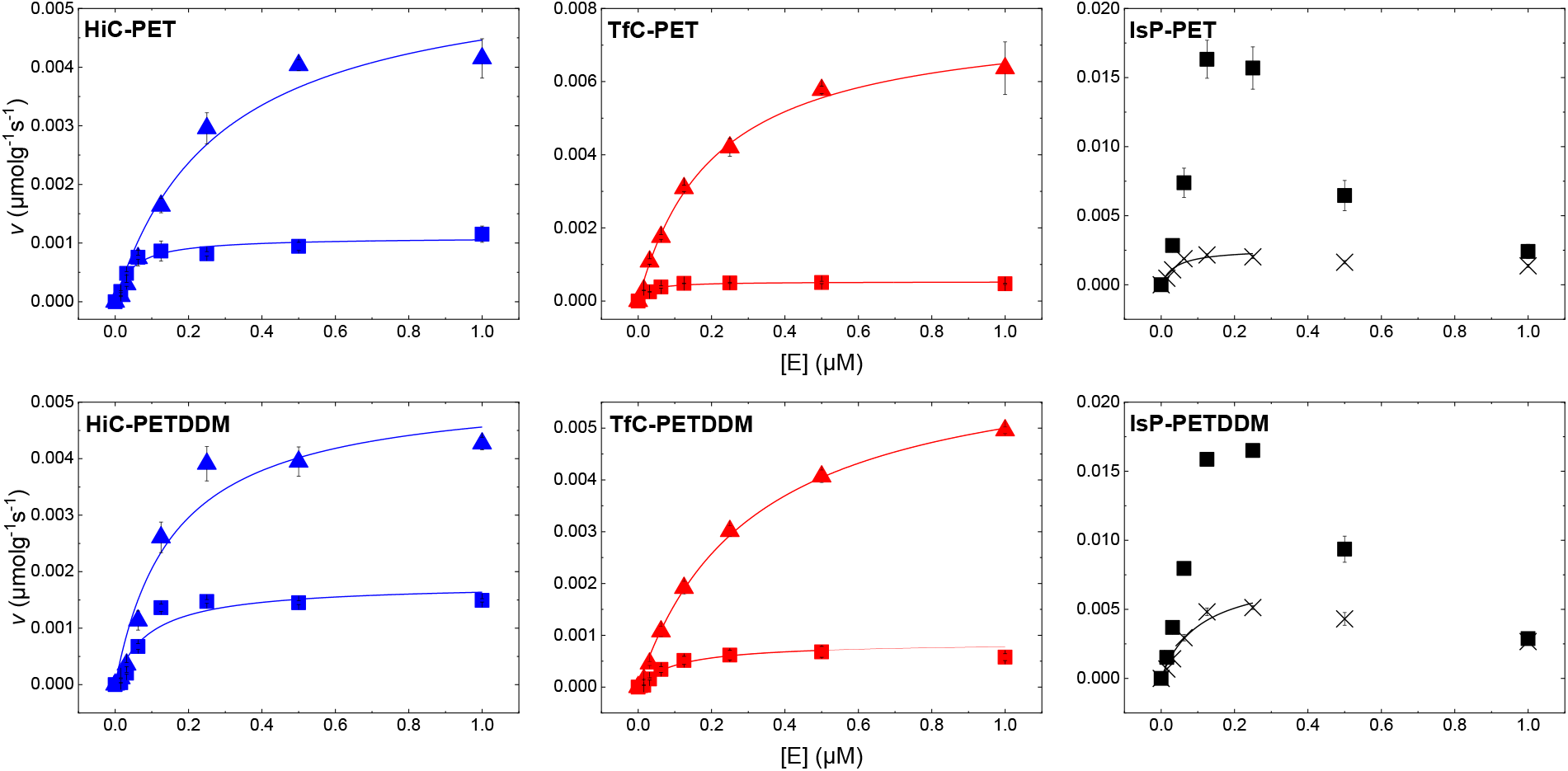
Inverse Michaelis Menten (^Inv^MM) plots showing the hydrolysis rate as a function of enzyme concentration. Symbols are experimental data from 3 h reactions at 40 °C (crosses), 50 °C (squares) and 60 °C (triangles) with 10 g/L PET. All symbols, colors and panel positions have the same meaning as in Fig. 3. Lines represent the best fit of eq. (5) and the derived kinetic parameters may be found in Tab. 2. IsP displayed a decline in activity upon high enzyme load and data for 50 °C were not possible to fit to eq. (5). IsP data for 40 °C, where this effect was less pronounced, were fitted until the drop.

**Table 1.** ^Conv^MM parameters determined for HiC, TfC and IsP on PET at 40, 50 or 60 °C with or without the addition of the nonionic surfactant DDM. It was not possible to resolve ^conv^K_M_ and k_cat_ from reactions at 40 °C with DDM for any of the enzymes. Numbers in brackets represent standard deviations of triplicate measurements.

| T (°C)/<br>Surfactant | HiC<br><sup>conv</sup>K<sub>M</sub><br>(gL<sup>-1</sup>) | k<sub>cat</sub>=<sup>conv</sup>V<sub>max</sub>/E<sub>0</sub><br>(s<sup>-1</sup>) | TfC<br><sup>conv</sup>K<sub>M</sub><br>(gL<sup>-1</sup>) | k<sub>cat</sub>=<sup>conv</sup>V<sub>max</sub>/E<sub>0</sub><br>(s<sup>-1</sup>) | IsP<br><sup>conv</sup>K<sub>M</sub><br>(gL<sup>-1</sup>) | k<sub>cat</sub>=<sup>conv</sup>V<sub>max</sub>/E<sub>0</sub><br>(s<sup>-1</sup>) |
| --- | --- | --- | --- | --- | --- | --- |
| 40/no | **ND\*** | **ND** | **ND** | **ND** | **4.9**<br>(± 0.90) | **0.52**<br>(± 0.053) |
| 50/no | **0.27**<br>(± 0.039) | **0.088**<br>(± 0.0021) | **1.2**<br>(± 1.1) | **0.015**<br>(± 0.0059) | **0.24**<br>(± 0.058) | **0.69**<br>(± 0.031) |
| 60/no | **0.26**<br>(± 0.16) | **0.043**<br>(± 0.0051) | **0.68**<br>(± 0.014) | **0.052**<br>(± 0.0034) | **ND** | **ND** |
| 50/DDM | **4.0**<br>(± 2.4) | **0.20**<br>(± 0.07) | **3.4**<br>(± 1.7) | **0.091**<br>(± 0.024) | **2.2**<br>(± 0.37) | **1.8**<br>(± 0.14) |
| 60/DDM | **4.7**<br>(± 3.0) | **0.12**<br>(± 0.043) | **1.2**<br>(± 0.65) | **0.081**<br>(± 0.015) | **ND** | **ND** |

\*ND: Not determined. Either due to experimental temperature above T<sub>m</sub> of the enzyme (IsP, 60 °C) or activity below experimental detection limit (HiC and TfC at 40 °C).

**Table 2.** ^Inv^MM parameters determined for HiC, TfC and IsP on PET at 40, 50 or 60 °C with or without the addition of the nonionic surfactant DDM. The parameters calculated for IsP are based on an approximated fitting of the data (see Fig. 4). Numbers in brackets represent standard deviations of triplicate measurements.

| T (°C)/<br>Surfactant | HiC<br>$^{inv}K_M$<br>($\mu M$) | $^{inv}V_{max}/S_0$<br>($\mu mol g^{-1} s^{-1}$) | TfC<br>$^{inv}K_M$<br>($\mu M$) | $^{inv}V_{max}/S_0$<br>($\mu mol g^{-1} s^{-1}$) | IsP<br>$^{inv}K_M$<br>($\mu M$) | $^{inv}V_{max}/S_0$<br>($\mu mol g^{-1} s^{-1}$) |
| --- | --- | --- | --- | --- | --- | --- |
| 40/no | ND* | ND | ND | ND | **0.039**<br>($\pm 0.017$) | **0.0026**<br>($\pm 3.5E-4$) |
| 50/no | **0.043**<br>($\pm 0.012$) | **0.0011**<br>($\pm 7.6E-5$) | **0.026**<br>($\pm 0.0067$) | **0.00053**<br>($\pm 2.5E-5$) | ND | ND |
| 60/no | **0.28**<br>($\pm 0.074$) | **0.0057**<br>($\pm 6.0E-4$) | **0.20**<br>($\pm 0.019$) | **0.0078**<br>($\pm 2.7E-4$) | ND | ND |
| 40/DDM | ND | ND | ND | ND | **0.11**<br>($\pm 0.038$) | **0.0078**<br>($\pm 0.0013$) |
| 50/DDM | **0.090**<br>($\pm 0.038$) | **0.0018**<br>($\pm 2.1E-4$) | **0.11**<br>($\pm 0.026$) | **0.00086**<br>($\pm 7.8E-5$) | ND | ND |
| 60/DDM | **0.16**<br>($\pm 0.052$) | **0.0053**<br>($\pm 5.9E-4$) | **0.31**<br>($\pm 0.025$) | **0.0065**<br>($\pm 2.2E-4$) | ND | ND |

\*ND: Not determined. Either not possible to fit data to the MM equation (IsP, 50 °C), experimental temperature above $T_m$ of the enzyme (IsP, 60 °C) or activity below experimental detection limit (HiC and TfC at 40 °C).

Some general trends in Tables 1 and 2 may be worth emphasizing. We found, for example, that well known efficacy of IsP against PET (37,38) relies on rapid turnover (k_cat_ is 1-2 orders of magnitude larger than for the two cutinases) rather than a particular affinity for the substrate (^conv^K_M_ values are comparable). Another trend in Table 1 is a conspicuous increase of both ^conv^K_M_ and k_cat_ upon the addition of a very low concentration of the nonionic surfactant DDM (50 μM, i.e. below the critical micelle concentration). Conversely, DDM had little effect on the parameters in the inverse regime.

In addition to the parameters in Tables 1 and 2, which result directly from non-linear regression, it is useful to consider some derived kinetic parameters. These include the attack site density, Γ_attack_, calculated according to eq. (3), and the catalytic efficacy (or specificity constant), ^mass^η=k_cat_/^conv^K_M_. The superscript of the specificity constant specifies that it is in mass based units, and we will discuss this further below (*c.f*. eq. (8)). Values of Γ_attack_ and ^mass^η for HiC, TfC and IsP acting on PET with or without DDM are listed in Table 3.

**Table 3.** The specificity constant (^mass^η) and the attack site density (Γ_attack_) for PET hydrolase reactions at 40, 50 and 60 °C with or without the addition of the surfactant DDM. Γ_attack_ is not reported for IsP due to the uncertain inverse maximal specific rates (see Table 2 and Fig. 4). Numbers in brackets represent standard deviations of triplicate measurements.

| T (°C)/<br>Surfactant | $^{\text{mass}}\eta$ (Lg<sup>-1</sup>s<sup>-1</sup>) | | | $\Gamma_{\text{attack}}$ (μmol/g) | |
| --- | --- | --- | --- | --- | --- |
|  | HiC | TfC | IsP | HiC | TfC |
| 40/no | **ND\*** | **ND** | **0.11**<br>(± 0.022) | **ND** | **ND** |
| 50/no | **0.33**<br>(± 0.048) | **0.013**<br>(± 0.013) | **2.9**<br>(± 0.71) | **0.013**<br>(± 0.018) | **0.035**<br>(± 0.014) |
| 60/no | **0.17**<br>(± 0.10) | **0.076**<br>(± 0.017) | **ND** | **0.13**<br>(± 0.090) | **0.15**<br>(± 0.019) |
| 40/DDM | **ND** | **ND** | **0.075**<br>(± 0.024) | **ND** | **ND** |
| 50/DDM | **0.050**<br>(± 0.035) | **0.027**<br>(± 0.015) | **0.82\*\***<br>(± 0.15) | **0.0090**<br>(± 0.0033) | **0.0095**<br>(± 0.0026) |
| 60/DDM | **0.026**<br>(± 0.019) | **0.068**<br>(± 0.039) | **ND** | **0.044**<br>(± 0.017) | **0.080**<br>(± 0.015) |

\*ND: Not determined. Either due to experimental temperature above $T_m$ of the enzyme (IsP, 60 °C) or activity below experimental detection limit (HiC and TfC at 40 °C).
\*\*Determined by linear fit, since not possible to extract the $^{\text{conv}}\text{MM}$ parameters $k_{\text{cat}}$ and $K_M$ from these data.

### Steady-state kinetics with BETEB as substrate

BETEB (Fig. 1b) may be seen as a fragment of a PET chain, and it has previously been used as a model substrate (21) for PET hydrolases. It is insoluble in water (equilibrium concentration in buffer could not be detected with the current methods), and we therefore used suspended substrate, and the same kinetic approaches as in the experiments with PET. We did not use surfactant in the experiments with BETEB. Products from BETEB hydrolysis were quantified by RP-HPLC, where peaks corresponding to the retention time of BA were dominant in all cases (Fig. S3). We hence used the built-up of BA to specify the steady-state rate. Like in the case of PET, we found that BsCE was essentially unable to hydrolyze BETEB. Results from ^conv^MM and ^inv^MM measurements for the three other enzymes and their analysis by eqs. (4) and (5) respectively are illustrated in Fig. 5.

**Figure 5.**
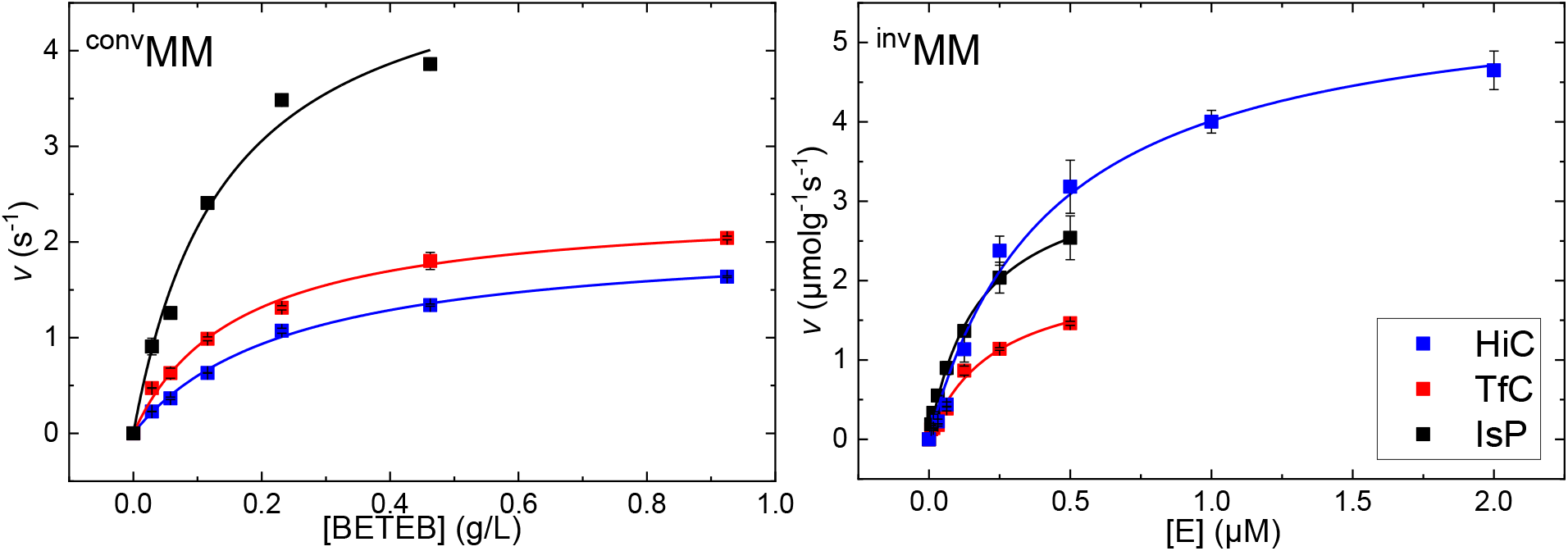
Conventional- and inverse MM plots for HiC (blue), TfC (red) and IsP (black), with initial hydrolysis rate as a function of BETEB or enzyme load. Symbols are experimental data from 10 min (IsP) or 20 min (HiC and TfC) reactions at 50 °C with 0.01 μM enzyme (^conv^MM) or 0.09 g/L BETEB (^inv^MM). Error bars represent standard deviations of duplicate measurements. Lines represent the best fit of the non-linear MM equation.

Kinetic parameters derived from Fig. 5 are listed in Table 4, and these data showed that HiC, TfC and IsP have values of ^conv^K_M_ that are similar and close to those observed with PET as substrate. In contrast to this similarity, turnover numbers on BETEB were orders of magnitudes larger than on PET. Finally, the density of attack sites on the BETEB surface was significantly higher than on PET.

**Table 4.** Kinetic parameters including Γ_attack_ and ^mass^η from conventional and inverse MM analyses on BETEB at 50 °C, calculated from the data in Fig. 5. Numbers in brackets represent standard deviations of duplicate measurements.

| **Enzyme** | **$^{\text{conv}}K_M$**<br>**($\text{gL}^{-1}$)** | **$^{\text{conv}}V_{\text{max}}/E_0$**<br>**($\text{s}^{-1}$)** | **$^{\text{inv}}K_M$**<br>**($\mu\text{M}$)** | **$^{\text{inv}}V_{\text{max}}/S_0$**<br>**($\mu\text{molg}^{-1}\text{s}^{-1}$)** | **$\Gamma_{\text{attack}}$**<br>**($\mu\text{mol/g}$)** | **$^{\text{mass}}\eta$**<br>**($\text{Lg}^{-1}\text{s}^{-1}$)** |
| --- | --- | --- | --- | --- | --- | --- |
| HiC | **0.25**<br>($\pm 0.021$) | **2.08**<br>($\pm 0.071$) | **0.43**<br>($\pm 0.064$) | **5.7**<br>($\pm 0.32$) | **2.7**<br>($\pm 0.18$) | **8.3**<br>($\pm 0.75$) |
| TfC | **0.16**<br>($\pm 0.019$) | **2.38**<br>($\pm 0.097$) | **0.23**<br>($\pm 0.045$) | **2.15**<br>($\pm 0.20$) | **0.90**<br>($\pm 0.15$) | **15**<br>($\pm 1.9$) |
| IsP | **0.14**<br>($\pm 0.029$) | **5.25**<br>($\pm 0.43$) | **0.17**<br>($\pm 0.012$) | **3.42**<br>($\pm 0.098$) | **0.65**<br>($\pm 0.057$) | **38**<br>($\pm 8.4$) |
| BsCE | **ND\*** | **ND** | **0.095**<br>($\pm 0.023$) | **0.15**<br>($\pm 0.012$) | **ND** | **ND** |

\*ND: Not determined, activities below experimental detection limit

### Steady-state kinetics on soluble substrates

Activity measurements of PET hydrolases on soluble substrates are convenient, but not generally indicative of the PET degrading capacity (5). Here, we included kinetic studies on *p*NP-val, MHET and BHET (see Fig. 1 for structures). Detailed data are presented in the Supporting information in the form of (conventional) MM plots (Fig. S4), and the calculated kinetic parameters K_M_ (mM) and k_cat_ (s^−1^) (Table S1). The specificity constants are presented in Table 5. The results show that HiC, TfC and BsCE hydrolyze *p*NP-val quite efficiently with η values between 2 × 10^4^ and 2 × 10^5^ M^−1^s^−1^ (k_cat_ values were between 4 s^−1^ and 30 s^−1^, Table S1). Table 5 also shows that none of the tested enzymes prefer MHET as substrate, but that all of them (particularly BsCE with η=2 × 10^4^ M^−1^s^−1^) are active on BHET. Finally, Table 5 confirms earlier reports (5,37), that IsP has very low activity against *p*NP-val.

**Table 5.** Specificity constants at 50 °C determined for HiC, TfC, IsP and BsCE on soluble substrates, including the PET fragments MHET and BHET as well as the model substrate *p*NP-val. Standard deviations of duplicate or triplicate measurements are shown in brackets. The underlying values of k_cat_ and K_M_ may be found in Table S1 of the Supporting information.

| Substrate/Enzyme | $\eta$ ($\text{M}^{-1}\text{s}^{-1}$) | | | |
| --- | --- | --- | --- | --- |
|  | HiC | TfC | IsP | BsCE |
| *p*NP-val | **1.90x10<sup>5</sup>**<br>($\pm 3.1 \times 10^4$) | **2.3x10<sup>4</sup>**<br>($\pm 3.6 \times 10^4$) | **4.4x10<sup>3</sup>**<br>($\pm 1.2 \times 10^3$) | **1.9x10<sup>4</sup>**<br>($\pm 2.1 \times 10^3$) |
| MHET | **ND\*** | **13**<br>($\pm 1.1$) | **ND** | **45**<br>($\pm 22$) |
| BHET | **5.5x10<sup>2</sup>**<br>($\pm 44$) | **1.9x10<sup>3</sup>**<br>($\pm 0.76 \times 10^3$) | **3.3x10<sup>3</sup>**<br>($\pm 2.1 \times 10^2$) | **1.7x10<sup>4</sup>**<br>($\pm 2.1 \times 10^3$) |

\*ND: Not determined, activities below the experimental detection limit.

## Discussion

Bioprocessing provides a promising tool in the fight against the immense environmental problems associated with an escalating consumption of plastic (7). The most progressed example is the use of enzymatic degradation of PET waste for recycling (4), but bioprocessing could also become important for remediation of microplastic pollution. This latter field has recently experienced important progress with the successful transformation of genes encoding plastic degrading enzymes into different microorganisms, which in turn becomes potential plastic scavengers (39–41). One common requirement for these applications is the design of better enzymes. This includes both better catalytic activity against the unnatural substrates and optimization with regards to relevant process conditions. Rational attempts to accomplish this will rely on a better understanding of structure-function relationships and formal kinetics makes up a key element in this respect. Nevertheless, formal kinetics with physically meaningful parameters remain scarce for plastic degradation and in the current work we propose and test a framework for this.

### Maximal turnover

The specific reaction rate at enzyme saturation, k_cat_=^conv^V_max_/E_0_, was determined for both soluble and insoluble substrates (Tables 1, 4 and S1). It appeared that the two cutinases HiC and TfC were quite slow on polymeric PET with k_cat_ values at 50 °C, corresponding to a few turnovers per minute (Table 1). IsP was significantly faster on PET with a k_cat_ of approximately 40 min^−1^ at 50 °C (and even 30 min^−1^ at 40 °C). These values may be compared with specific rates from some earlier reports. Tournier et al. (2020) included TfC (BTA hydrolase 1 in their terminology) in an extensive study of PET hydrolases (4). They used amorphous PET particles of a similar size as the current, and reported specific rates corresponding to respectively 0.02 s^−1^ (50 °C) and 0.07 s^−1^ (60 °C). These values compare very well with k_cat_ for TfC in Table 1. Tournier et al. did not report K_M_ values so we cannot say whether these specific rates reflect enzyme saturation, but the substrate load was 2 g/L, which is above K_M_ values found here, so the results probably represent reasonable estimates of k_cat_. Barth et al. (2015), on the other hand, found a much higher turnover number of about 2 s^−1^ at 60 °C for a related cutinase from *T. fusca* (13). This latter work used PET nanoparticles (about 100 nm) as substrate, and the higher turnover calls for further investigations of relationships between maximal turnover and particle size.

We consistently observed much higher k_cat_ values on the model substrate BETEB. Specifically, the three enzymes with activity on PET (HiC, TfC and IsP) showed quite similar k_cat_ on BETEB in the range of 2-5 s^−1^ at 50 °C (Table 4). For TfC, this is two orders of magnitude faster than its turnover number on PET, and for HiC and IsP it is one order of magnitude faster. It is interesting to notice that this difference in k_cat_ occurred although both PET and BETEB are insoluble, and this obviously suggests that the interfacial nature of the reaction does not *per se* dictate a slow turnover. This interpretation is further supported by comparisons with k_cat_ values for the same enzymes acting on the soluble PET fragments MHET and BHET. These latter turnover numbers (Table S1) were comparable to those found on BETEB, and again contradicts any direct correlation between insoluble substrate and slow turnover. It is also of interest to compare the observed k_cat_ values with typical values for enzymes acting on their preferred, natural substrate. For esterases, a survey of the BRENDA database suggested that k_cat_ values for native, soluble substrates predominantly fell in the 3 - 30 s^−1^ range (42), and other meta-analyses covering wider selections of enzymes have found similar average k_cat_ values (43,44). We note that the k_cat_ values found here for different PET fragments (whether soluble or insoluble) were in this range. This may be unexpected for HiC and TfC, which (unlike IsP) are probably devoid of any evolutionary adaptation to the substrate. Overall, these observations indicate that the turnover rate of intact PET depends on interactions in the polymeric substrate, whereas the chemical steps associated with the hydrolytic reaction (which are common to PET and its fragments) are comparably fast. Thus, interactions between polymer strands in the solid substrate could result in large activation barriers for complexation or dissociation, and hence slow down the overall process, even if the actual hydrolytic reaction is fast (as indicated by k_cat_ values for PET fragments). Analogous arguments have previously been put forward in discussions of the (slow) enzymatic hydrolysis of cellulose (45,46).

### Specificity constants

The overall efficacy of enzymes (natural or engineered) that act on anthropogenic substrates is typically gauged by the specificity constant (47). For insoluble substrates this parameter is readily calculated according to eq. (6)

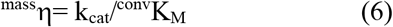

The superscript designates that this definition leads to specificity constants in mass-based units (in the current case (g/L)^−1^ s^−1^), and the application of ^mass^η defined in this way, is essentially limited to comparisons of isoenzymes acting on the same substrate. Here, we found that ^mass^η for IsP (≈ 3 s^−^ 1(g/L)^−1^ at 50 °C) was much higher than ^mass^η for HiC and TfC, and this again testifies the superior performance of IsP on PET. The specificity constants on BETEB were higher and quite similar for the three enzymes (between 10 and 40 s^−1^(g/L)^−1^ at 50 °C). In other words, IsP appeared distinctly superior to the two cutinases on polymeric PET, but not on the shorter BETEB substrate. For a broader discussion of specificity constants, we converted the ^mass^η values to the conventional units of M^−1^s^−1^. Specifically, we used the attack site density, Γ_attack_, from Table 3 to calculate the Michaelis constant, ^conv^K_M_, in molar units, (*i.e.* moles of attack sites per liter suspension at half saturation)

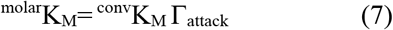

Combining eqs. (6) and (7) yields an expression for the molar specificity constant, ^molar^η

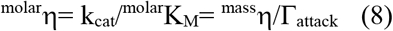

Specificity constants for the insoluble substrates were calculated according to eq. (8) and are listed in Table 6. We used the symbol ^molar^η to indicate that these values were derived indirectly. Nevertheless, we will discuss them together with specificity constants for soluble substrates (denoted η in Table 5) calculated in the normal way.

**Table 6.**
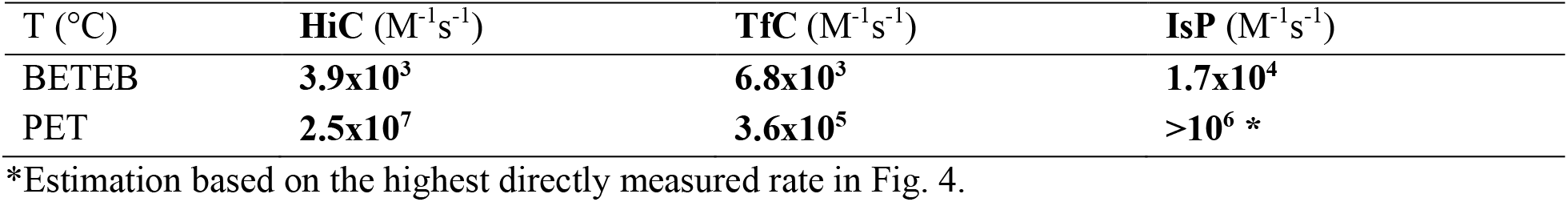
Molar specificity constants (^molar^η) calculated according to eq. (8) for HiC, TfC and IsP acting on the insoluble substrates PET and BETEB at 50 °C (without the addition of DDM).

| T (°C) | HiC ($\text{M}^{-1}\text{s}^{-1}$) | TfC ($\text{M}^{-1}\text{s}^{-1}$) | IsP ($\text{M}^{-1}\text{s}^{-1}$) |
| --- | --- | --- | --- |
| BETEB | **$3.9 \times 10^3$** | **$6.8 \times 10^3$** | **$1.7 \times 10^4$** |
| PET | **$2.5 \times 10^7$** | **$3.6 \times 10^5$** | **$>10^6$ \*** |

\*Estimation based on the highest directly measured rate in Fig. 4.

If we first consider soluble substrates, we found very low specificity constants on MHET for all investigated enzymes, while the values on BHET were in the range from 10^3^ to 10^4^ M^−1^s^−1^. To put this into perspective, we note that the majority of enzymes acting on their preferred, natural substrate have specificity constants of 10^4^ to 10^6^ M^−1^s^−1^ (44). The smaller specificity constants for soluble PET fragments indicates that these are poor substrates for most of the enzymes investigated here, even if k_cat_ is fairly high (see above). The only clear exception for this is BHET hydrolysis by BsCE, which showed a specificity constant (2 × 10^4^ M^−1^s^−1^) comparable to (the low end of) natural enzyme-substrate systems. Interestingly, ^molar^η-values for insoluble substrates were larger than η for the soluble PET fragments. Thus, for HiC and TfC, molarη attained values of 10^7^ and 10^5^ M^−1^s^−1^, respectively, on polymeric PET at 50 °C. Unfortunately, the value for IsP could not be determined due to the problems of finding the inverse maximal rate (see Fig. 4), but estimates based on the highest directly measured rate in Fig. 4, suggest that ^molar^η is at least 10^6^ M^−1^s^−1^ for IsP on PET. High specificity constants for the degradation of soluble, synthetic compounds have been observed before (see (47) for a review), but it is noteworthy that this parameter increased for insoluble substrates. While the exact meaning of molarη remains to be elucidated, one possible explanation is that the enzyme adsorbs nonspecifically on the hydrophobic surface of the substrate particles. If indeed so, enzymes will concentrate near the attack sites and hence experience a higher effective substrate concentration compared to bulk reactions. In essence, this means that the reaction space is reduced from 3D to 2D. This interpretation finds some support in the observation of very strong, non-specific adsorption of cutinases on other types of (non-hydrolysable) plastic (48,49). It is also worth noting that similar ^molar^η values have been reported earlier for other interfacial enzyme reactions. Specifically, hydrolysis of (insoluble) microcrystalline cellulose by the cellulases Cel6A and Cel7A showed ^molar^η in the range of 10^5^ to 10^6^ M^−1^s^−1^ (50,51). These cellulases have separate carbohydrate binding modules (CBM), which promotes strong surface adsorption, and this again suggests a link between adsorption and high ^molar^η of interfacial reactions. Interestingly, CBMs also show affinity for PET surfaces, and fusion proteins with a cutinase and a CBM have shown improved activity against PET (52,53). Possible relationships between adsorption and the specificity constant may be further illustrated by considering the two terms in eq. (8) (k_cat_ and ^molar^K_M_) separately. As discussed above, k_cat_ for HiC and TfC on PET were much lower than typical values for enzymes modifying their innate substrate, and it follows that the high values of ^molar^η rely on an unusually low ^molar^K_M_. To illustrate this, we inserted data for HiC and TfC acting on PET at 50 °C into eq. (7). This gave ^molar^K_M_ values of 30-40 nM for both enzymes, and it follows that the hydrolytic rate reaches half its maximal value, when there are some 40 nmol attack sites per liter suspension. This is a sign of a very strong substrate interaction. Thus, K_M_ for wild type enzymes catalyzing naturally occurring reactions in the bulk phase predominantly fall in the range between 10 μM and 1 mM (44). We propose that this anomalously low K_M_ could be mediated by nonspecific adsorption to the PET surface, which promotes the encounter of enzyme and substrate.

### Attack site densities and effects of surfactant

Many earlier studies of enzymatic PET degradation have adopted the use of surfactants in the protocols. This has served different purposes including modification of enzyme-substrate interactions, cleaning of the PET surface or simply experimental convenience in the preparation and handling of PET suspensions (14,23,52,54–56). To systematically test the effect of a nonionic surfactant, we repeated both the conventional- and inverse MM measurements on the PET substrate in a buffer that was supplemented with DDM. We found that even a low surfactant concentration (ca. 50 μM), imparted strong and systematic effects on the ^conv^MM parameters (Table 1), while parameters from the ^inv^MM measurements (Table 2) were only marginally affected. Specifically, ^conv^K_M_ and k_cat_ increased in the presence DDM, and this meant that we could not always reach saturation in the ^conv^MM measurements (in particular the ^conv^MM curve for IsP at 40 °C, was almost linear, see lower right panel in Fig. 3). Therefore, the conventional kinetic parameters with DDM are only approximate, but the increase was distinctive (Fig. 3). The effect of the surfactant also emerged as a marked reduction of the number of attack sites recognized by the enzymes (Table 3). These observations may bring some clues of the underpinning mechanisms. Thus, the results are in line with the interpretation that DDM accumulates on the hydrophobic PET surface, and hence screen a fraction of the attack sites. This may explain both the lowered Γ_attack_, and the increased ^conv^K_M_, as a higher mass-load of PET would be required to reach half-saturation if some attack sites are covered. Interestingly, negative effects of surface coverage are compensated by an increase in k_cat_, and the overall picture is that DDM leads to fewer, but more rapidly converted complexes. Molecular origins of this compensation remains to be investigated, but it could be related to unproductive binding of enzyme. Thus, if a fraction of the enzymes adsorbs nonproductively (or with poor productivity), k_cat_ will reflect a weighted average of active and inactive populations. Nonionic surfactants are known to reduce nonspecific adsorption, and this could diminish putative populations of nonproductively bound enzyme and hence raise the observed k_cat_. At any rate, the pronounced effects of DDM call for more systematic investigations of this area both with respect to molecular mechanism and potential significance for bioprocessing of polyester. Some important work using either anionic- or cationic surfactants has pointed towards changed electrostatic interactions at the interface as the origin of improved PET hydrolase efficacy (23,55), but the current observations suggest that even nonionic surfactants may modify the kinetics distinctively, and hence that other factors are relevant too.

### Conclusions

The understanding of enzyme-catalyzed hydrolysis of PET and other plastics is not complete and there is no well-established framework for kinetic analyses of the reaction. Here, we have tested an approach to this problem, which is based on the introduction of putative attack sites on the PET surface. We showed how the number of attack sites can be readily determined experimentally and used to convert the substrate mass load into an apparent molar concentration of catalytically competent sites in the suspension. This opened up for the use of mass-action kinetics, and the introduction of kinetic parameters for comparative analyses. We conducted such analyses for four enzymes, and identified distinctive differences in substrate affinity, turnover rates, catalytic efficiency and the ability to locate attack sites on PET. We also demonstrated that the approach opens for kinetic comparisons of the catalytic performance on respectively intact PET and smaller (soluble or insoluble) fragments of the polymer.

Kinetics makes up the experimental link between the structure and function of catalysts (57), and we propose that the approach presented here may become a useful tool within PET hydrolase enzymology. This is both with regards to discussions of molecular mechanisms, rate limiting steps and rational design of enzymes with improved activity against this man-made substrate.

## Experimental procedures

### Enzymes

Two cutinases, HiC [AAE13316.1] from *Humicola insolens* and TfC [AAZ54921.1] from *Thermobifida fusca* were expressed and purified as described previously (58,59). The carboxyl-esterase, BsCE [P37967.2] from *Bacillus subtilis* was expressed as secreted proteins in *B. subtilis* and purified as described previously (21). IsP, a His-tagged variant (S238F/W159H) of the PETase from *Ideonella sakaiensis* [6EQD_A] was expressed as secreted proteins in *B. subtilis* and purified in two steps by Ni-affinity chromatography followed by gel filtration on a HiLoad 26/600 Superdex 75 pg column. The molar enzyme concentration was determined by Abs280 and the calculated extinction coefficient (60) for the respective enzyme. The thermal transition midpoint (T_m_) of the enzymes was determined by differential scanning fluorimetry using a Prometeus NT.48 instrument (Nano Temper, Munich, Germany). Enzyme samples (in 50 mM acetate buffer pH 5.0) with concentrations of approximately 0.5 mg/mL were heated from 20 to 95 °C at a rate of 200 °C/h.

### Substrates

Semi-crystalline PET powder (Product number ES306030) was purchased from Goodfellow Co (UK). The crystallinity reported by the producer was >48%. Particle sizes determined by laser diffraction ranged from 10-500 μm with a dominance of sizes around 200μm (see Fig. S1, Supporting information). Terephtalic acid (TPA), bis(2-hydroxyethyl) terephthalate (BHET) and *p*-nitrophenyl valerate (*p*NP-val), used as substrates and/or standard samples for spectrophotometric and reversed-phase high-performance liquid chromatography (RP-HPLC) analysis, were purchased from Sigma-Aldrich. Mono(2-hydroxyethyl) terephthalate (MHET), used as standard sample and substrate, was produced by enzymatic hydrolysis of BHET (5 hours contact time with HiC). The oligomeric PET model substrate bis(2-(benzoyloxy)ethyl) terephthalate (BETEB) was synthesized from BHET (25 g, 98 mmol) and benzoyl chloride (28.5 mL, 245 mmol) in pyridine (100 mL). BHET was dissolved in pyridine and cooled on ice-water. The flask was equipped with a pressure-equalizing addition funnel, magnetic stirring and nitrogen. Benzoyl chloride was added dropwise and the mixture was left stirring at room temperature overnight. Heavy precipitation made it necessary to add more pyridine (20 mL). For work up, DCM (250 mL) and ice-water (500 mL) was added and the mixture transferred to a separation funnel. The aqueous phase was extracted with additional DCM (200 mL). The combined DCM-phases were washed with 0.1 M HCl (2 * 250 mL) and sat. NaHCO_3_ (2 * 200 mL), then dried (Na_2_SO_4_), filtered and evaporated. The crude product was recrystallized twice from warm anhydrous EtOH (250 mL). The yield was 41 g (90%) off-white solid. An overview of the chemical structures of the substrates may be found in Fig. 1. We note that PET and BETEB are essentially insoluble in buffer and investigated as stirred suspensions (see below). The other substrates, *p*NP-val, BHET and MHET, were soluble over the concentration ranges used here. All enzyme assays were conducted in 50 mM sodium phosphate buffer, pH 8.0 (except the *p*NP-val assay, see below). The PET suspension was prepared either with the addition of 0.0025% (w/V) of the nonionic surfactant, n-dodecyl β-D-maltoside (DDM; Sigma-Aldrich) or without any surfactant.

### Activity assay with *p*NP-val

Enzyme-mediated *p*-nitrophenol (*p*NP) release from *p*NP-val was detected continuously over 10 min at 405 nm. Kinetic measurements were performed in 96-well plates using a plate reader (Molecular Devices SpectraMax Paradigm). Reactions contained 150 μL *p*NP-val and 30 μL enzyme dissolved in 50 mM TRIS-HCl buffer, pH 7.7, with concentrations of *p*NP-val ranging from 0-0.83 mM. Measurements were performed in triplicates at 25 °C. Linear regression of a standard curve obtained with known concentrations of *p*NP was used for quantification. Data were fitted to the MM equation using ORIGIN PRO 2019 (OriginLab Coorporation, Northhampton, MA, USA).

### Activity assay with PET

#### ^Conv^MM analysis

A suspension-based plate reader assay (32) was adapted for initial rate measurements of PET degrading enzymes. Reactions were performed in triplicates in 250 μL volumes with 0.03 μM enzyme and PET loads from 0-6 g/L, using low binding microplates (Greiner Bio-One™ 655900). The plates were sealed and kept at the selected temperature (40 °C, 50 °C or 60 °C) in an incubator/shaker (KS 4000 ic control, operated at 450 rpm, IKA, Staufen, Germany). The contact time was 2 hours for IsP and 5 hours for HiC and TfC. These contact times were selected to get a good signal from the relatively slow enzyme reaction without exceeding the linear part of the progress curve (see Fig. S2). The reactions were stopped by centrifugation (3 min, 3500 rpm) and 100 μL of supernatant was transferred to UV-transparent microplates (Corning) for spectrophotometric measurements in a plate reader at 240 nm. Enzymatic product formation was quantified against standard curves of BHET. As described elsewhere (32) this standardization provides a reasonable measure of the overall activity. Finally, data were fitted to the MM equation using ORIGIN PRO 2019.

#### ^Inv^MM analysis

Assay conditions, product quantification and data processing for ^inv^MM analysis were similar to the procedures for ^conv^MM described above, except that the PET load was 10 g/L, and the enzyme concentration varied between 0-1 μM. The plates were kept at the experimental temperature in the incubator/shaker for 3 hours and it was confirmed that this corresponded to the linear part of the progress curve (Fig. S2).

### Activity assay with BETEB

Enzyme-catalyzed BETEB hydrolysis was performed in a similar manner as the PET reactions, but analyzed by RP-HPLC due to the high background absorbance of BETEB in plate reader analysis. Samples with BETEB suspensions were incubated in an Eppendorf thermomixer at 50 °C, 1100 rpm between 10 and 40 minutes, depending on enzyme. Reactions for ^conv^MM analysis were performed with 0.01 μM enzyme and BETEB loads between 0 and 0.92 g/L. Reactions for ^inv^MM analysis used 0.092 mg/mL BETEB and enzyme concentrations from 0 to 2 μM. After incubation, the samples were centrifuged and 100 μL supernatant was redrawn. HCl was added to the supernatant and treated samples were stored in the freezer to reduce auto hydrolysis prior to RP-HPLC analysis. Enzymatically produced benzoic acid (BA, Fig. 1g) was quantified against standard curves of BA, and data were fitted to the MM equation using ORIGIN PRO 2019.

### Activity assay with soluble PET fragments

Activity on the PET fragments MHET and BHET was also assayed in low binding microplates. Reactions were performed in duplicates in 250 μL volumes with substrate concentrations from 0-2 mM. Reactions were incubated in an Eppendorf thermomixer at 50 °C, 1100 rpm between 10 min and 2 hours, depending on enzyme and substrate. Enzyme concentrations were 0.1 μM (Except for HiC on BHET: 0.5 μM and BsCE on BHET: 0.005 μM). All reactions were stopped by addition of HCl and stored in the freezer prior to RP-HPLC analysis. Enzymatically produced MHET and TPA were quantified against standard curves of the same compounds, and data were fitted to the MM equation using ORIGIN PRO 2019.

### Reaction products detected by RP-HPLC

The concentration of TPA, MHET and BA from selected enzyme reactions with BETEB, BHET and MHET was determined by RP-HPLC (Chemstation series 1100, Hewlett Packard). The instrument was equipped with a diode array detector and an ODS-L optimal column from Capital HPLC (25 × 4.6 mm) packed with C18 particles 5 μm in diameter size. Injection volume was 20 μL and samples were eluted with 24% acetonitrile over 25 minutes. Products were identified based on absorption at 240 nm. Flow rate was set to 0.5 mL/min and the column was kept at 40 °C. Peak analysis was performed using the ChemStation for LC 3D software. Standards with known concentrations of TPA, MHET, BHET and BA were used to quantify reaction products for kinetic analyses. Duplicates and substrate blanks (for quantification of auto hydrolysis) were included for all reactions.

## Data availability

All data that support the findings of this study are included in the published article and its Supporting information file.

## Acknowledgements

We would like to thank Anni Bygvrå Hougaard and Linda Rikke Dietrich, University of Copenhagen and Claus Sternberg, DTU-Bioengineering’s Bioimaging Platform for assistance in the characterization of PET particle size.

## Abbreviations

BA: Benzoic acid
BETEB: Bis(2-(benzoyloxy)ethyl) terephthalate
BHET: Bis(2-hydroxyethyl) terephthalate
DDM: n-Dodecyl β-D-maltoside
EG: Ethylene glycol
Γ_attack_: density of attack sites
MHET: Mono(2-hydroxyethyl) terephthalate
MM: Michaelis-Menten
PET: Poly(ethylene terephthalate)
*p*NP-val: *p*-Nitrophenyl valerate
RP-HPLC: Reversed-phase high-performance liquid chromatography
TPA: Terephthalic acid

